# Metabolomic profiling reveals grade-specific niacinamide accumulation and its therapeutic Potential via SIRT1-CD38-EMT axis modulation in cervical cancer progression

**DOI:** 10.1101/2024.12.17.628866

**Authors:** Shivani Jaiswal, Vivek Mishra, Srija Majumder, Pramod P. Wangikar, Shinjinee Sengupta

## Abstract

Despite the availability of new therapies for cervical cancer, innovative strategies are essential to address challenges related to drug resistance and high toxicity. The present study focuses on the metabolic profiling of cervical carcinoma using non-targeted metabolomics approach using liquid chromatography-mass spectrometry. Our study identified over 100 metabolites in cervical tissue samples (both cancerous and adjacent normal) using HILIC and reversed-phase chromatography in the positive and negative ionization modes. The major metabolic alterations included changes in glycolysis, the citric acid cycle, amino acid metabolism, nucleotide metabolism, and nicotinamide metabolism in a grade-dependent manner. Interestingly, metabolic differences between HPV-positive and HPV-negative tumors included elevated arginine and proline metabolism, nicotinamide metabolism, and phosphatidylcholine biosynthesis. We further validated our findings by analyzing transcriptomics datasets from the Gene Expression Omnibus database to understand the expression patterns of the underlying genes involved in the dysregulated pathways. We observed that nicotinamide metabolism exhibits significant effects in lower-grade cervical cancers and specific HPV genotypes. We treated cervical cancer cell lines with niacinamide (NAM), an amide form of niacin, to evaluate its potential therapeutic efficacy. NAM treatment modulated NAD+ metabolism with key players such as CD38, PARP, NAMPT, and SIRT1, promoting apoptosis and inhibiting cell proliferation in cervical cancer cells. Importantly, the metabolic responses differed between the HPV-positive SiHa cells and HPV-negative C33A cells, reflecting distinct NAD+ metabolic adaptations. The study highlights the metabolic adaptation or shifts in cancer progression and provides insights into NAM’s molecular mechanisms and therapeutic potential for precision medicine in cervical cancer.

## 1.1. Introduction

Cervical carcinoma (CC) exhibits poor prognosis due to distant recurrence and metastasis[1]. Previous studies have established the fact that cervical cancer is associated with human papillomavirus (HPV) infection resulting in p53 degradation[2],[3]. Cervical cancer predominantly spreads through lymphatic vessels and direct extension[4]. Primary treatment options include radical hysterectomy, radiotherapy, and cisplatin-based chemotherapy[5]. However, careful consideration of specific treatment schemes is necessary based on the cancer types and HPV status. Despite these interventions, more than 30% of patients exhibit radio resistance, and the recurrence rate of early-stage cervical cancer exceeds 5% within 4.5 years[6]. Additionally, the standard-of-care radiation therapy for locally advanced cervical cancer has remained unchanged for over three decades, resulting in stagnant clinical outcomes[7]. Consequently, there is an urgent need for the development of novel treatment strategies to enhance therapeutic efficacy.

Since the 1920s, Otto Warburg, known as “Warburg effect,” made a groundbreaking observation regarding the increase in the glucose consumption and lactate production in tumors compared to normal tissues[8]. In the presence of oxygen, the cancer cells for energy production prefer glycolysis over oxidative phosphorylation[9]. Over time, our understanding of cancer-associated metabolic changes has expanded, leading to the recognition of metabolic reprogramming as a hallmark of cancer [10], [11]. While most cancers exhibit multiple hallmarks, different cancer types and even subtypes within the same cancer can display diverse metabolic phenotypes[12]. The metabolic traits of cervical cancer cells have also garnered increasing attention in recent studies, offering novel insights for diagnosis and treatment[13], [14]. Recent studies have highlighted the potential of metabolic remodeling to effectively augment cervical cancer therapy by improving chemo-and radiosensitivity[15].

Cancer significantly impacts metabolism by reprogramming cellular metabolic pathways and modifying metabolites level, these aberrant metabolites play vital role in metastasis and hold potential to serve as a biomarker for personalized cancer therapy[16],[17]. In this report, we have characterized the metabolic profiling of cervical cancer patient biopsies using a nontargeted metabolic profiling strategy based on liquid chromatography-mass spectrometry which offers tremendous potential for clinical oncology by enabling the identification of cancer-related metabolites. We identified more than 100 different metabolites from the tissue metabolome in each category grade wise as well as in presence and absence of HPV infection. The primary metabolic aberrations in tumors tissues when compared to normal included changes in glycolysis, the citric acid cycle, amino acid metabolism, nucleotide metabolism, and niacinamide (nicotinamide) metabolism. Further, niacinamide (NAM) treatment in cervical cancer cell lines, such as C33A and SiHa, modulated NAD^+^ metabolism by targeting key players like CD38, PARP, NAMPT, and SIRT1, inducing apoptosis and reducing proliferation, likely through effects on the EMT pathway. The distinct metabolic responses between HPV-positive and HPV-negative cell lines underscore the complexity of NAD^+^ metabolism in cervical cancer. Thus, the study aims to assess targeted biochemical pathways linked to cervical cancer with a focus on high-resolution analysis. The validation of findings is carried out using relevant cell lines, strengthens the translational relevance of the results.

## 2. Results

### 2.1. Clinical characteristics

Several previous studies have highlighted the accumulation or depletion of several metabolites leasing to changes in the metabolic profiling of cervical patents[14],[18],[13]. In this study we examine the metabolic changes in a grade dependent and HPV occurrence in cervical tumor biopsies and further validate the predicted biomarker for early detection in cervical cancer cells. For the current study, we used a small cohort of cervical human cancer biopsies from Indian individuals with different grades (**Sup Table 1).** We initially verified the cancerous nature of these tissues by employing hematoxylin and eosin (H&E) staining (**Figure 1a**). The findings revealed varied H&E staining patterns, alongside the presence of neoplastic cells exhibiting intense nuclear staining, suggestive of hyperproliferative cell presence. We isolated the genomic DNA to check the presence of HPV in the biopsy samples (**Fig. S1a**). Western blot analysis revealed degraded p53 or total absence of p53 protein specially in the HPV positive biopsies (**Fig. S1b**). It is a well-known fact that p53 is not in detectable expression in HPV positive tissue as it readily degraded by HPV. The absence of lower expression of p53 in HPV negative tissues could be due to frameshift or truncated mutation[19], or negative regulation by MDM2[20]. Further, to understand the metabolic changes, tumor biopsies were grouped into three categories, such as Grade 2, Grade 3 and normal tissues (**Figure 1b**). For each group 4 or more patient tumor biopsies were taken. We further categorized the entire cohort as HPV positive or HPV Negative biopsies. We observed that major portion of the changes occurred in the amino acid metabolisms including their derivatives in our cohort of cervical tumor tissues when compared the adjacent normal tissues (**Figure 1c**). Majority of the lipids of the class PC/LPC was observed to be dysregulated in the overall data (**Figure 1c**).

**Figure 1.**
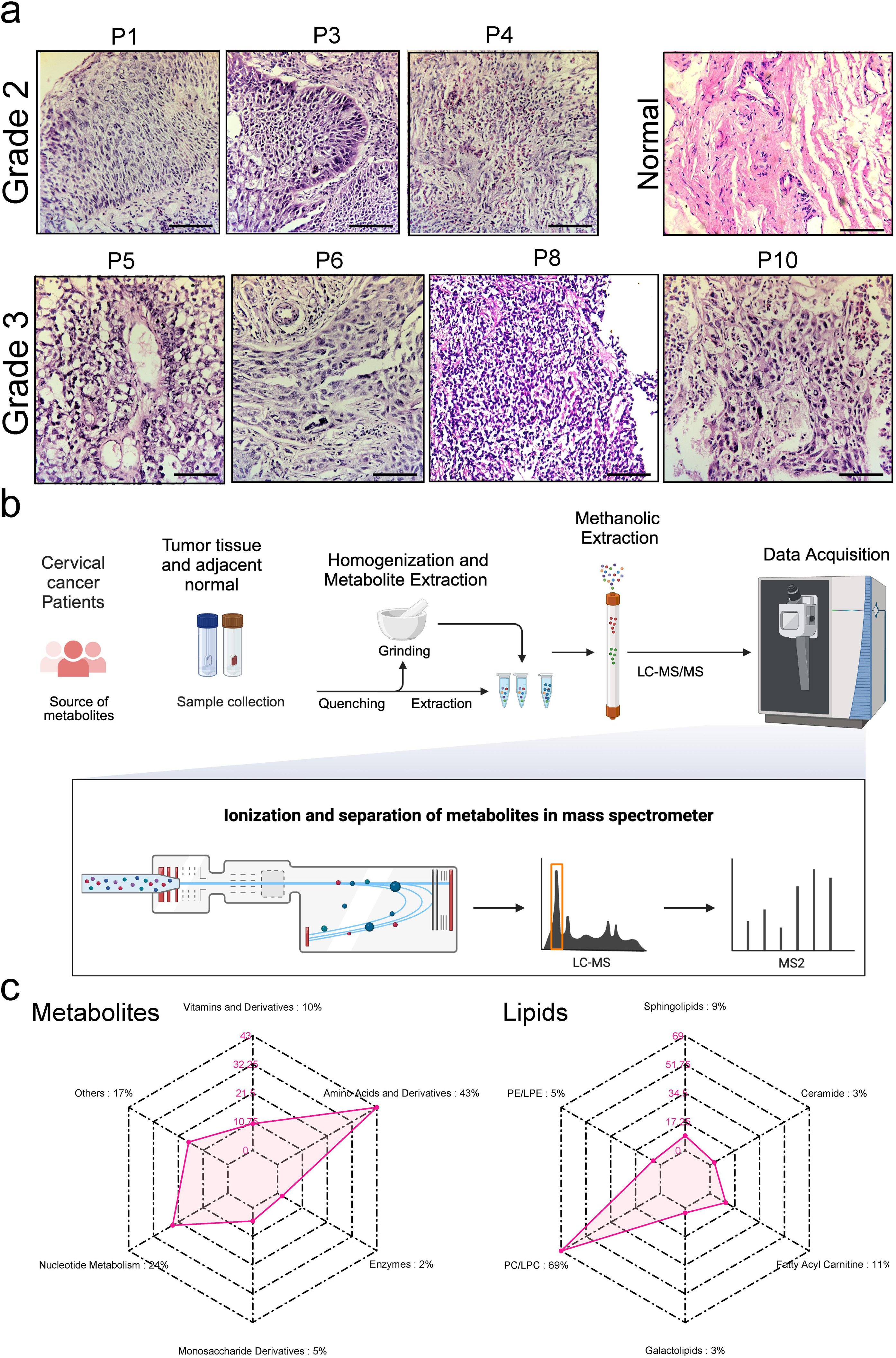
Characterization of cervical cancer cohort and schematic workflow of metabolic analysis. a. Hematoxylin and eosin staining of cervical cancer tissue sections from different patient biopsies. The tissue sections were deparaffinized, hydrated, and stained with hematoxylin and eosin before being examined under a microscope at 40x magnification. Representative images of cervical cancer tissues of varying grades are highlighted with black boxes along with normal tissue for comparison. These images exhibit differential staining patterns with hematoxylin, which intensely stains the numerous nuclei present. The scale bars represent 30 μm. The images are representative of three independent experiments. b. The schematic diagram outlines the experimental workflow, including sample collection from cervical cancer patients, tumor biopsies, tissue lysis using bead beater homogenizer, metabolite extraction, to obtain the methanolic extract for metabolite and lipid analysis in LC-MS/MS followed by statistical analysis for identifying biomarkers. c. Radar plot (Left) representing total metabolites and lipids (Right) dysregulated in different categories which may be altered in cervical cancer tissues of our cohort when compared to the adjacent normal tissues. The below chart shows the top dysregulated lipids falling in different categories across normal and tumor cervical tissue samples, indicating potential changes in lipid metabolism.

### 2.2. Metabolic profiling of grade wise cervical cancer biopsies

Metabolic profiling displayed distinct profile separation between cancer and adjacent normal tissues as demonstrated by PLS-DA plot (**Figure 2a**). Furthermore, when plotted separately, there was also a clear segregation between healthy individuals and the two cancer grades (**Figure 2b)**. 17 metabolites showed a statistically different fold change ≥ ±1.3 in normal vs. all cervical cancer tissues (**Figure 2c**). These metabolites are adenosine diphosphate (ADP), adenosine 3’-monophosphate (3’AMP), dimethylarginine, 2-Phenylacetamide, tyrosine, phosphoethanolamine, choline, glutamic acid, urocanic acid, timonacic, proline, choline phosphate, N6-Isopentenyladenine, niacinamide, trimethoprim (TMP), hypoxanthine, 1-Myristoyl-2-palmitoyl-sn-glycero-3-phosphocholine (MPPC) and adenosine-5’-diphosphate Di(monocyclohexylammonium) salt.

**Figure 2.**
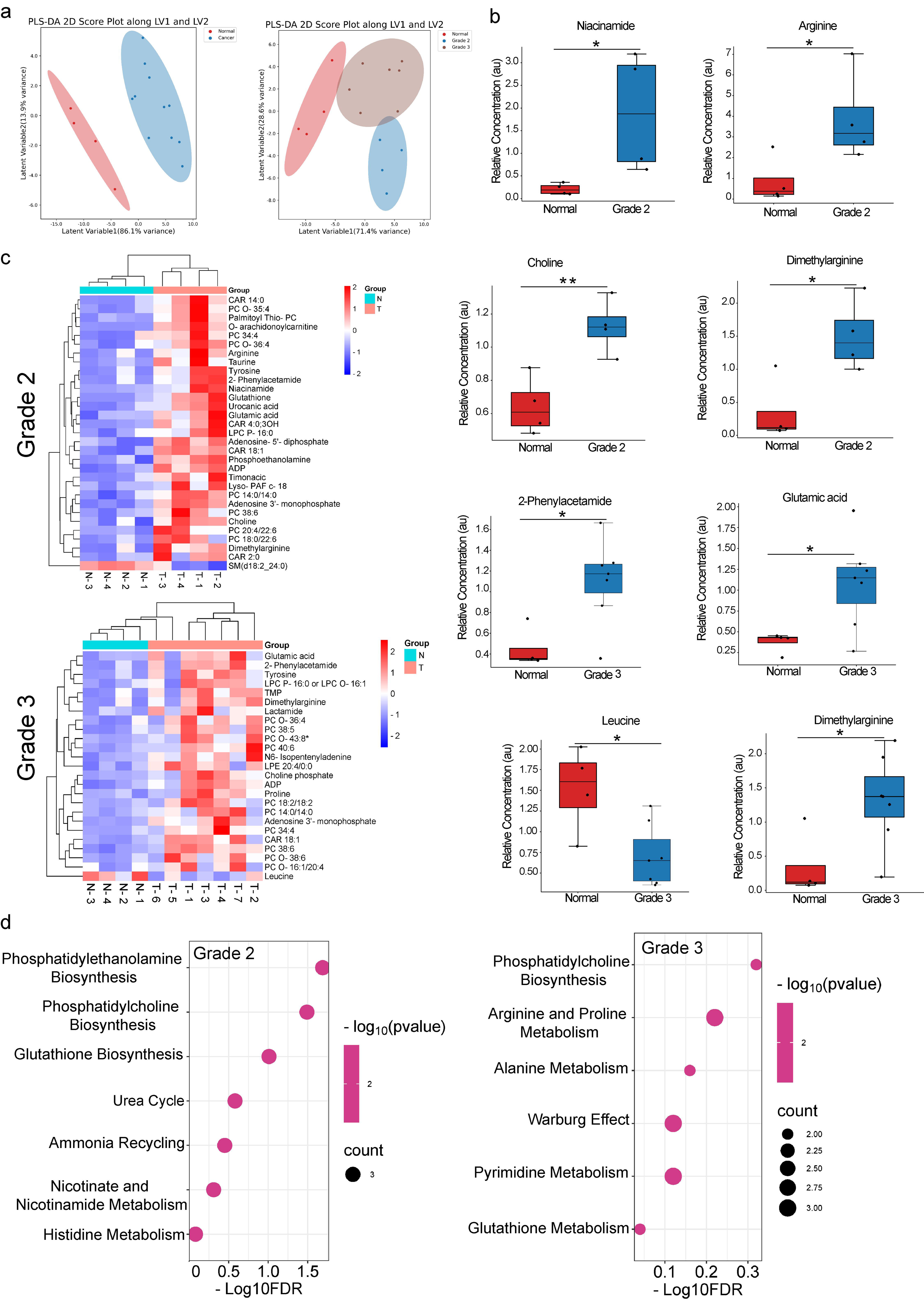
Metabolic changes in grade 2 and grade 3 cervical tumor biopsies. The figure presents metabolomic data comparing various metabolite levels across normal, grade 2, and grade 3 cervical tissue samples. a) Partial least squares discriminant analysis (PLS-DA) 2D score plot showing the separation of normal and cancer (left) and Normal versus grade 2, 3 cancer samples (right) based on their metabolic profiles along the latent variables LV1 and LV2. Each point represents an individual sample, and the shaded areas represent 95% confidence intervals. b) Bar plots showing the relative concentration levels of niacinamide, arginine, dimethylarginine, and choline in normal and grade 2 cancer tissue samples. * indicates statistically significant difference between groups (p < 0.05). Bar plots depicting the relative concentration levels of 2-phenylacetamide, leucine, and dimethylarginine in normal and grade 3 cancer tissue samples. * indicates statistically significant difference between groups (p < 0.05). c) Heatmap showing the relative concentration levels of significant metabolites in normal and grade 2 (upper panel) and normal and grade 3 (Lower panel) cancer tissue samples. Rows represent individual samples, and columns represent different metabolites. The color scale ranges from blue (low concentration) to red (high concentration). d) Enrichment bubble plot with respect to reference library of Metaboanlyst showing overall significant enriched metabolite sets. For each group 4 or more patient tumor biopsies were taken.

Analyzing the cervical cancer patients when divided into high- and low-grade indicates that 13 metabolites showed a statistically significant fold change ≥ ±1.3 when low grade patient tissues were compared with adjacent normal tissues. These were 3’AMP, ADP, O-arachidonoylcarnitine, phosphoethanolamine, glutathione, urocanic acid, palmitoyl thio-PC, timonacic, dimethylarginine, tyrosine, taurine, 2-Phenylacetamide, niacinamide and arginine (**Figure 2b, c**). On comparison of higher-grade patient tissues with adjacent normal tissues, only 11 metabolites displayed statistically significant fold change such as ADP, adenosine 3’-monophosphate, leucine, 2-Phenylacetamide, N6-Isopentenyladenine, tyrosine, TMP, choline phosphate, dimethylarginine, glutamic acid and lactamide. The dysregulated metabolites shown are involved in various important enriched sets that are crucial for cell survival, proliferation and signaling (**Figure 2d**). Comparison of tissues between Grade 2 and Grade 3 mostly, revealed dysregulation of lipids such as phosphatidylcholines, and sphingomyelins (**Fig. S2a**). Clear segregation was observed among the different histological grades in PCA plot (**Fig. S3a**).

We observed that several of these lipids were upregulated in the tumor biopsies. CAR 18:1 is significantly upregulated in grade 2 and grade 3 cancer samples compared to adjacent normal tissue (**Figure 3**). Increased levels of fatty acids like CAR 18:1 can provide cancer cells with energy and building blocks for membrane synthesis, supporting rapid proliferation and tumor growth.

**Figure 3.**
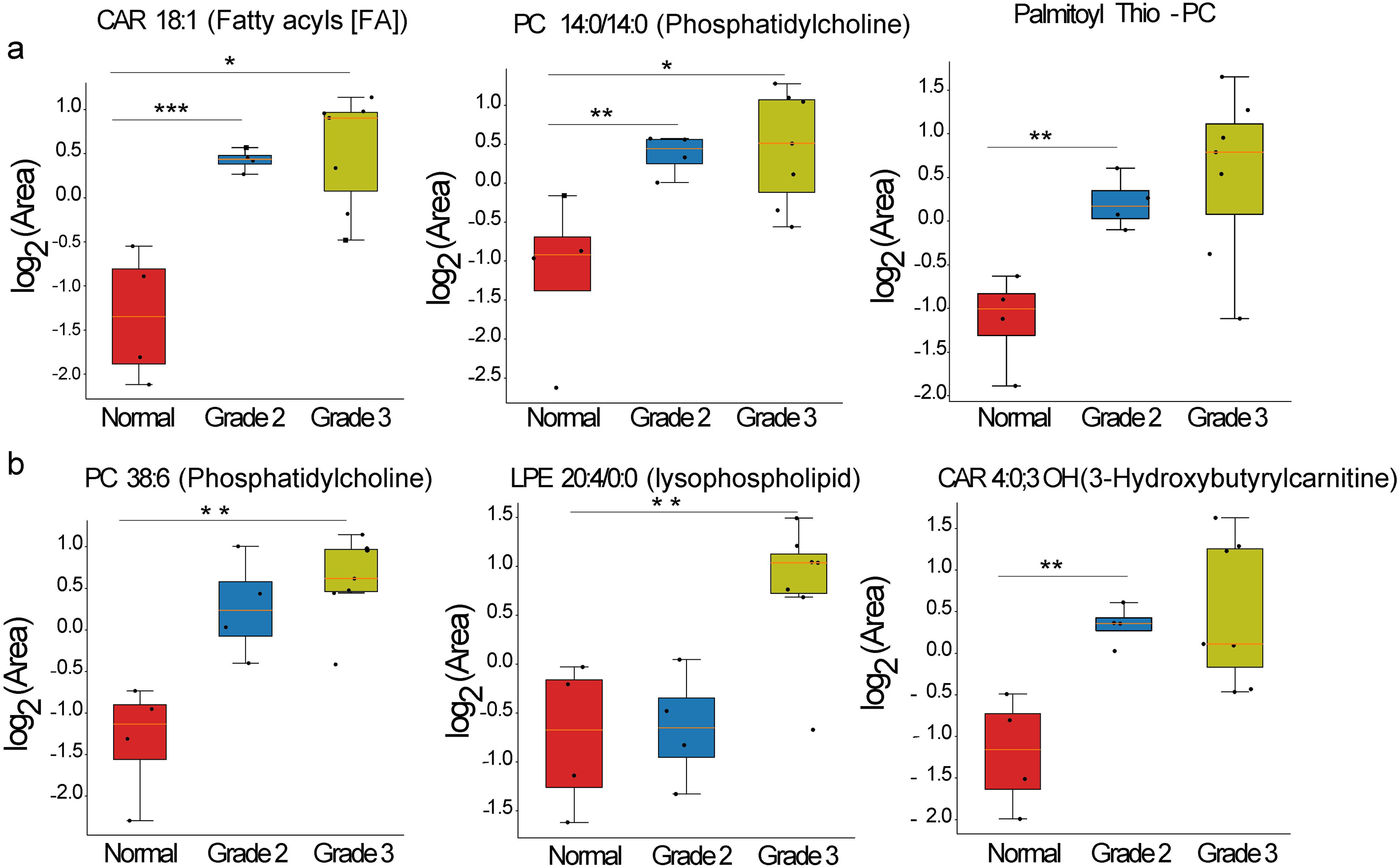
Changes in the lipid accumulation in grade wise cervical patient biopsies. The figure shows changes in the levels of various lipid metabolites across normal, grade 2, and grade 3 cervical tissue samples. a) The graphs represent the log2 area values of CAR 18:1 (Fatty acyls [FA]), PC 14:0/14:0 (Phosphatidylcholine), and Palmitoyl Thio-PC, respectively, between the different tissue groups. Asterisks indicate statistically significant differences. b) The plots depict the log2 fold changes in the levels of PC 38:6 (Phosphatidylcholine), LPE 20:4/0:0 (lysophospholipid), and CAR 4:0/3-CH/3-Hydroxybutyrylcarnitine across the tissue groups. CAR refers to acylcarnitines, which are fatty acid derivatives involved in the transport of fatty acids into the mitochondria for β-oxidation. PC and LPE represent phosphatidylcholine and lysophosphatidylethanolamine, respectively, which are major lipid components of cell membranes and play roles in membrane structure and signaling.

Similarly, PC 14:0/14:0 (Phosphatidylcholine) and PC 38:6 (Phosphatidylcholine) show elevated levels in higher cancer grades. Phosphatidylcholines are major components of cell membranes, and their upregulation can facilitate the formation of new membrane structures required for rapid cell division and proliferation in cancer cells[21]. Palmitoyl Thio-PC derivative is significantly increased in grade 3 cancer samples, since it is like other phosphatidylcholines, it can contribute to membrane biogenesis and support the high proliferative rate of cancer cells. LPE 20:4/0:0 lysophospholipids was observed to accumulate in grade 3 cancers as they can promote cancer progression by modulating signaling pathways involved in cell growth, migration, and invasion, as well as contributing to membrane remodeling and lipid metabolism in cancer cells. CAR 4:0/3 CH/3-Hydroxybutyrylcarnitine is elevated in grade 3 cancer samples possibly to its role in the transport of fatty acids into mitochondria for energy production, which is essential for the high metabolic demands of rapidly dividing cancer cells.

### 2.3. Metabolic profiling of presence and absence of HPV in cervical cancer biopsies

We performed PCR to segregate the tumor samples based on HPV infection (**Fig. S1a**). These groups were analyzed to study the differences in the metabolic alterations. Significant differences in the levels of niacinamide, xanthine, choline, and proline between normal and HPV-positive samples were observed, suggesting alterations in metabolic pathways related to these metabolites in HPV-associated cancers (**Figure 4a**). The heatmap demonstrate distinct metabolic profiles between normal and HPV-positive samples, with several metabolic pathways showing enrichment or depletion in HPV-positive samples and in HPV-positive samples compared to normal (**Figure 4c, d**). The metabolic pathways most enriched in HPV-positive samples include arginine and proline metabolism, nicotinate and nicotinamide metabolism, phosphatidylcholine biosynthesis, and phosphatidylethanolamine biosynthesis, among others. Upregulation of lipids such as PC 38:6 (a phosphatidylcholine species) and CAR 18:1 (a fatty acyl) in HPV-positive samples, suggesting their alterations in lipid metabolism pathways in HPV-associated cancers (**Figure 4d**). Notable lipid species such as SM 41:10/02 and PC O-43:8, exhibited differential expression in HPV positive tissues when compared to HPV negative tissues (**Fig. S2b**). The Partial Least Squares Discriminant Analysis (PLS-DA) 2D score plot shows clustering based on latent variables (LV1 and LV2) among three groups, Normal, HPV Negative (HPVNeg), and HPV Positive (HPVPos) (**Figure 4e**). The clustering indicates a clear separation of metabolic profiles between the three groups. Normal samples cluster distinctly from both HPV-negative and HPV-positive groups, while HPV-positive and HPV-negative samples share some partial overlap, suggesting differences in metabolic composition across these groups. PCA plot also highlights the distribution of HPV status across the samples, with partial overlap between the HPV positive and negative groups (**Fig.S3b**). Pathway analysis in HPV-positive tissues shows enrichment in Phosphatidylethanolamine Biosynthesis and Phosphatidylcholine Biosynthesis, indicating altered phospholipid metabolism. Other enriched pathways include Nicotinate and Nicotinamide Metabolism and Arginine and Proline Metabolism. These suggest a shift in energy and redox metabolism in HPV-positive samples, possibly to support cancerous growth and maintain cellular redox balance (**Figure 4f, Top panel**). Enriched pathways in HPV-negative samples include Purine Metabolism and Histidine Metabolism, which may indicate a focus on nucleotide synthesis and amino acid metabolism. Other pathways such as Phenylalanine, Tyrosine, Tryptophan Biosynthesis, Nitrogen Metabolism, and Phenylalanine Metabolism are also highlighted, suggesting differences in amino acid and nitrogen metabolism compared to HPV-positive samples (**Figure 4f, Below panel**). Overall, the data suggests that HPV infection is associated with distinct metabolic reprogramming, particularly in pathways related to lipid metabolism, nucleotide synthesis, and energy production, potentially contributing to the proliferative and survival advantages of HPV-positive cancer cells.

**Figure 4.**
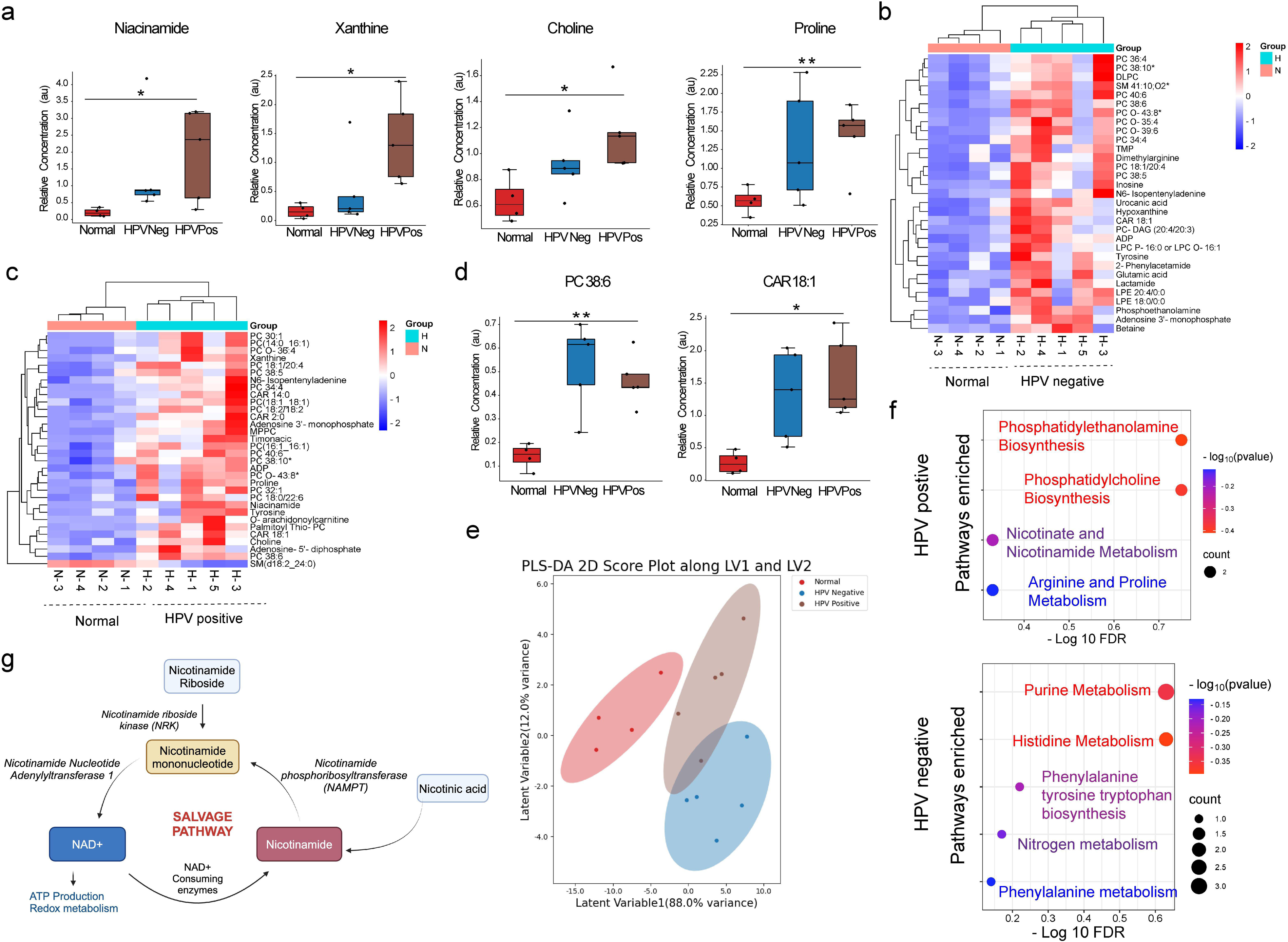
Metabolic changes in HPV positive and negative patients. a) Bar plots showing the relative concentrations of some of the significant metabolites such as niacinamide, xanthine, choline, and proline in normal, HPV-negative (HPV Neg), and HPV-positive (HPV Pos) tissue samples. * indicates a statistically significant difference (p < 0.05) compared to adjacent normal. b) Heatmap visualization of the relative significant metabolite concentrations with metabolite names listed on the right for patients biopsies with HPV negative and normal tissues. c) Heatmap visualization of the relative significant metabolite concentrations with metabolite names listed on the right for patients biopsies with HPV positive and normal tissues. d) Bar plots showing the relative concentrations some of the lipids such as PC 38:6 and CAR 18:1 in normal, HPV-negative, and HPV-positive samples. * indicates a statistically significant difference (p < 0.05) compared to adjacent normal. e) Partial least squares discriminant analysis (PLS-DA) 2D score plot showing the separation of normal with HPV positive and negative cancer samples based on their metabolic profiles along the latent variables LV1 and LV2. Each point represents an individual sample, and the shaded areas represent 95% confidence intervals. f) Enrichment analysis based on the HPV positive and negative patents. g) Schematic showing salvage pathway and role of NAM.

### 2.4. Metabolic pathway analysis using transcriptomics GEO datasets

To obtain an overview of the underlying genes affecting these metabolic pathways, we retrieved microarray data from NCBI (Gene Expression Omnibus: GSE63514) and performed correlational analysis with our metabolomics data (**Figure 6 a-e**). For metabolic pathway analysis using transcriptomics GEO datasets, we selected genes commonly involved in metabolic reprogramming of our respective enriched pathways from the Gene Set Enrichment Analysis (GSEA) Human Gene Set data. When we overlapped the transcriptomics data with the GSEA data, we observed that several genes affecting the major metabolic pathways were dysregulated, aligning with our metabolomics findings. Transcriptomics data exhibited metabolic pathway alterations in accordance with our data. We observed alteration in glutathione metabolism, which was also evident as GCLC shows upregulation, while GGT6 and GPX3 are downregulated, suggesting alterations in cellular redox balance. For Warburg effect genes such as PIK3CA, SLC2A1, and VEFA are upregulated, consistent with increased glycolysis in cancer cells. The upregulation of PIK3CA strongly supports the Warburg effect by promoting glycolytic metabolism and growth signaling (**Figure 6a,b**). SLC2A1 encodes GLUT1, a glucose transporter that facilitates the entry of glucose into cells. Our data exhibited alteration in arginine and proline metabolism which is quite evident as ADC1 and GAM1 show downregulation, while MAOB is upregulated. Similarly, sphingolipid metabolism was shown affected due to significant upregulation of UGT8, while SMPD2 is downregulated, indicating changes in membrane lipid composition.

Interestingly, we observed a notable upregulation of CD38 in tumors samples, suggesting an increased capacity for NAD^+^ generation and adenosine production potentially affecting the nicotinate and nicotinamide pathway (**Figure 6 c,d**). Further we checked for the expression and survival analysis of CD38 and NT5E (also known as CD73) genes across the cervical cancer (CESC) patients via Box plot and Kaplan-Meier survival plot using GEPIA2 as these genes affect the nicotinamide pathway (**Figure 6e**). These changes collectively point to a shift in NAD^+^ metabolism that may support the increased energy demands of more aggressive tumors.

### 2.5. Nicotinamide as a potential biomarker

The effects of nicotinate and nicotinamide metabolism were observed in both grade wise category as well as due to HPV genotype (**Figure 5**). Thus, we validated the effect of the metabolite in HPV positive and negative cervical cancer cell lines to enhance its translational relevance. To directly demonstrate the effect of niacinamide on cervical cancer cells, we treated HPV positive SiHa and HPV negative C33A cells with different concentrations of NAM. IC_50_ of NAM was calculated with SiHa and C33A cell line as 36.36 mM and 40.01 mM respectively (**Fig. S2)**. To analyse the CD38 expression, we treated C33A and SiHa cells with 40.1 mM and 36.6 mM NAM respectively. qPCR analysis revealed downregulation of CD38 in both cell lines. In SiHa, NAMPT was upregulated, while its levels remained unchanged in C33A. Additionally, PARP expression was downregulated in SiHa but upregulated in C33A cells (**Figure 6f**). qPCR analysis also revealed that Tfam (mitochondrial transcription factor A) was upregulated in SiHa cells following NAM treatment. FACS analysis revealed ∼25% of cell apoptosis (both early and late combined) due to niacinamide treatment in SiHA cells compared to control cells (**Figure 7a**). The effect was more pronounced in HPV negative cells, where ∼37% of cell population exhibited apoptosis (both early and late combined). This observance was also in accordance with the increase in the expression of apoptotic marker BAX (**Figure 7b**). To understand the mechanism behind NAM mediated apoptosis, we studied the expression of some of the epithelial-mesenchymal transition (EMT) markers. Vimentin levels were reduced in C33A cells after 24h of NAM treatment compared to untreated controls suggesting NAM may inhibit EMT and metastatic potential (**Figure 7b**). Similar effect was also observed in other EMT markers like beta-catenin; however, the effect was more pronounced in SiHa cells than C33A. Furthermore, TGF-β was downregulated in both cell lines after NAM treatment, indicating a potential mechanism through which NAM affects EMT (**Figure 7c**). NAM treatment also upregulated an NAD^+^-dependent deacetylase, SIRT1 (Sirtuin 1) in SiHa cells. In addition, NAM treatment reduced phosphorylated AKT (p-AKT) levels in both cell lines, while total AKT levels remained unchanged. The expression of c-Myc was also downregulated in both SiHa and C33A cells, further supporting NAM’s role in inhibiting pro-survival pathways and promoting apoptosis (**Figure 7 b,c**). Similar observation was observed for wound healing assay, where the distance traveled by cells to close an artificial wound is measured. The data shows that NAM treatment for 24h significantly reduces the distance in both SiHa and C33A cells compared to untreated controls at 0h and 24h, suggesting NAM inhibits cell proliferation and wound healing capacity (**Figure 7 d,e**). Further a reduction in colony forming unit (CFU) absorbance was observed for both the cell lines treated with nicotinamide (**Figure 7f**). This suggests that nicotinamide treatment may have inhibited cell growth or proliferation compared to untreated controls. Additionally, immunofluorescence analysis showed reduced expression of the proliferative marker, Ki67 in treated SiHa and C33A cells compared to untreated cells (**Figure 7g**). The co-localization with DAPI staining indicated a decrease in cell proliferation in the treated groups, further confirming the impact of treatment on cellular growth dynamics. Thus, the data confirms the therapeutic potential of niacinamide especially in both HPV positive and negative cervical cancers.

**Figure 5.**
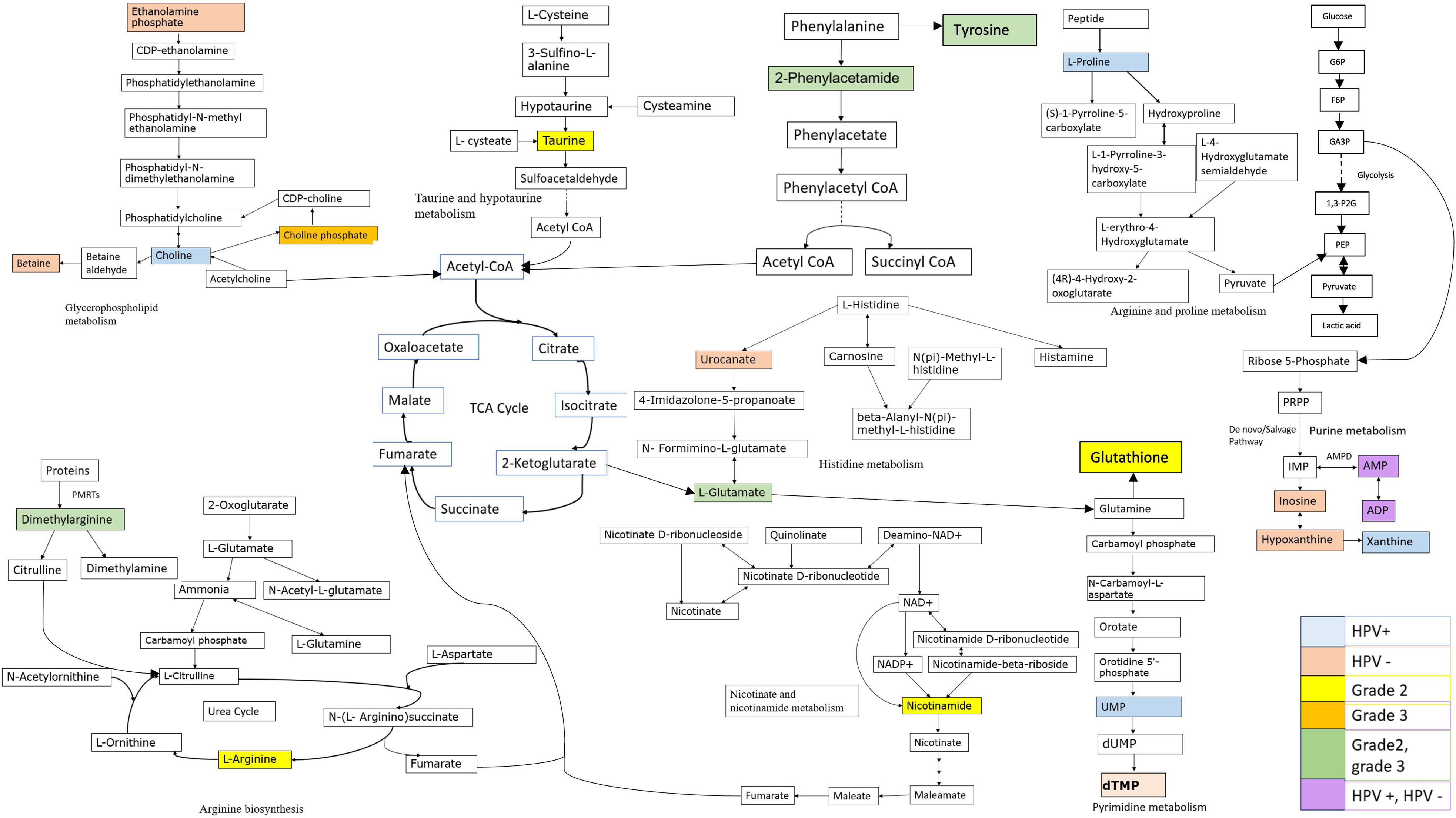
Overall pathway analysis based on differential metabolites. The figure depicts various metabolic pathways and highlights several metabolites that are dysregulated in cervical cancer patients. The pathways shown include glycolysis, the citric acid cycle, amino acid metabolism (including phenylalanine, tyrosine, histidine, and arginine metabolism), and nucleotide metabolism. The schematic also highlights various enzymes and metabolic intermediates involved in these pathways, providing a comprehensive overview of the metabolic alterations observed in cervical cancer patients. Several metabolites seen dysregulated such as nicotinamide, a precursor of NAD^+^, which plays a crucial role in energy metabolism and is often dysregulated in cancer cells. Others like glutathione which is an important antioxidant molecule involved in maintaining cellular redox homeostasis, which can be dysregulated in cancer cells, Inosine, nucleoside involved in purine metabolism, which can be altered in cancer cells due to increased nucleotide synthesis demands. The color coding shows the category in which the metabolite was differentially present.

## 3.0 Discussion

Gene expression and cellular regulation often culminate in the production of metabolites, which are essential indicators of physiological and pathological processes [22]. The process of tumor formation can potentially alter the overall metabolism within the human body[23]. This is evident from the distinct metabolic profiles observed in biofluids, such as plasma, urine, and other bodily fluids, of individuals diagnosed with various cancers[24]. Our study provides insight into metabolic changes during cervical cancer progression, with a specific focus on HPV-induced carcinogenesis and grade-dependent alterations. These metabolic changes offer potential therapeutic targets and biomarkers for early detection.

Our metabolic profiling identified over 100 dysregulated metabolites in tumor tissues, including key metabolites involved in nucleic acid synthesis, lipid metabolism, and energy production (**Figure 2**). Niacinamide and arginine play critical roles in nucleic acid synthesis, crucial for rapid cancer cell division[25],[26], while arginine also contributes to nitric oxide signaling and protein methylation[27]. Changes in lipid metabolites, such as phosphatidylcholine and lysophospholipids, point to alterations in membrane composition and energy metabolism, essential for tumor cell survival and proliferation[28],[29] (**Figure 3**). These alterations may reflect the cancer cells’ increased demand for energy and structural components, particularly in higher-grade tumors[30]. The observed changes in metabolite levels across the tissue groups suggest alterations in various metabolic pathways, potentially associated with the progression of cervical cancer.

In terms of energy metabolism, our results aligned with previously reported transcriptomic datasets, showing the upregulation of ADP and glutathione, reflecting high glycolytic turnover and oxidative stress response (**Figure 6**). The upregulation of glutathione observed in our study could be a protective response by tumor cells against oxidative stress, stemming from their rapid metabolic activity. We noted a complex regulation of glutathione-related genes, suggesting a potential alteration in the tumor’s ability to manage oxidative stress. The upregulation of niacinamide is indicative of enhanced NAD^+^ synthesis, essential for glycolysis and metabolic pathways, which align with the Warburg effect seen in advanced cervical cancer. The metabolic reprogramming observed in our study encompasses changes in energy production, lipid metabolism, and redox balance, which are critical for tumor survival and progression.

**Figure 6.**
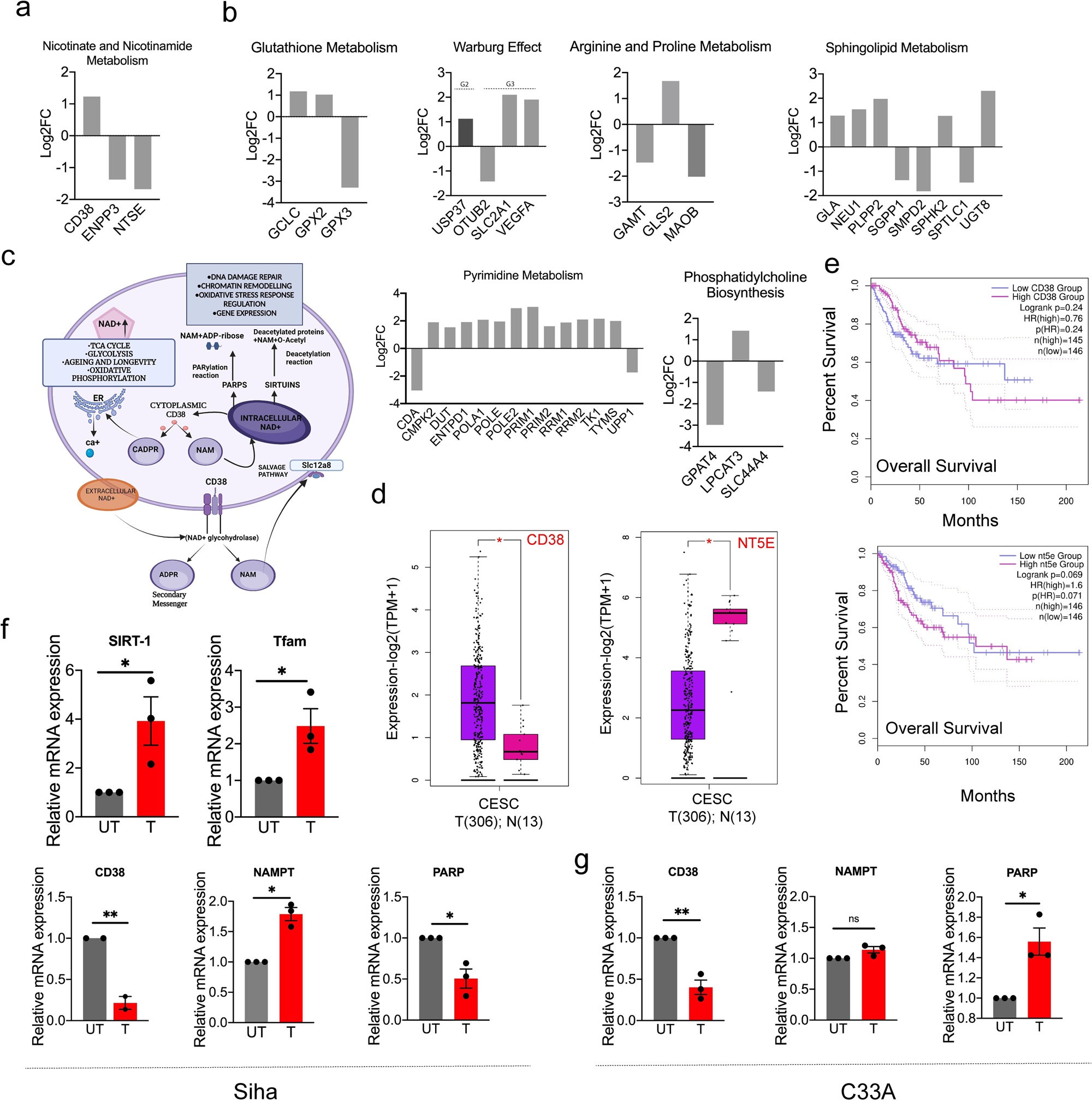
Analysis of dysregulated metabolic pathway using transcriptomics GEO datasets. a. Nicotinate and Nicotinamide pathway: CD38 is upregulated as this gene are involved in NAD^+^ metabolism and salvage pathways. The y-axis represents the log2 fold change in gene expression, with positive values indicating upregulation in tumors and negative values indicating downregulation compared to normal. b. Glutathione Metabolism: GPX3 is downregulated, while GCLC is upregulated in tumors, suggesting alterations in cellular redox balance. Warburg effect: Most glycolytic enzymes are upregulated in tumor samples consistent with increased aerobic glycolysis. Arginine and Proline Metabolism: MAOB is Upregulated, while others are downregulated indicating changes in amino acid metabolism. Additional pathway alterations such as Sphingolipid Metabolism: Sphingolipid-related genes such as SMPD2 is downregulated and rest all upregulated. Similarly, genes underlying the pyrimidine metabolism and phophophatidylcholine biosynthesis pathways are shown. c. Schematic diagram of the Nicotinate and Nicotinamide pathway, highlighting CD38’s role in NAD^+^ metabolism and various cellular processes. d. Box plots comparing the expression of CD38 and NT5E in cervical squamous cell carcinoma (CESC) tumor samples (T, n=306) versus normal samples (N, n=13). Asterisks indicate statistical significance. e. Kaplan-Meier survival curves showing overall survival of patients based on CD38 and NT5E expression levels. Left: CD38 expression; Right: NT5E expression. High and low expression groups are compared, with log-rank p-values and hazard ratios (HR) provided. This figure demonstrates the complex interplay between metabolic pathways, gene expression, and patient outcomes in cancer, with a particular emphasis on CD38’s role in these processes. f. qPCR showing upregulation of Sirt-1 and Tfam, downregulation of CD38 in SiHa cell lines on NAM treatment. NAMPT is upregulated in SiHa while PARP was downregulated in SiHa. g. qPCR showing downregulation of CD38 in C33A cell lines on NAM treatment. NAMPT level remains unchanged while PARP was upregulated in SiHa.

**Figure 7.**
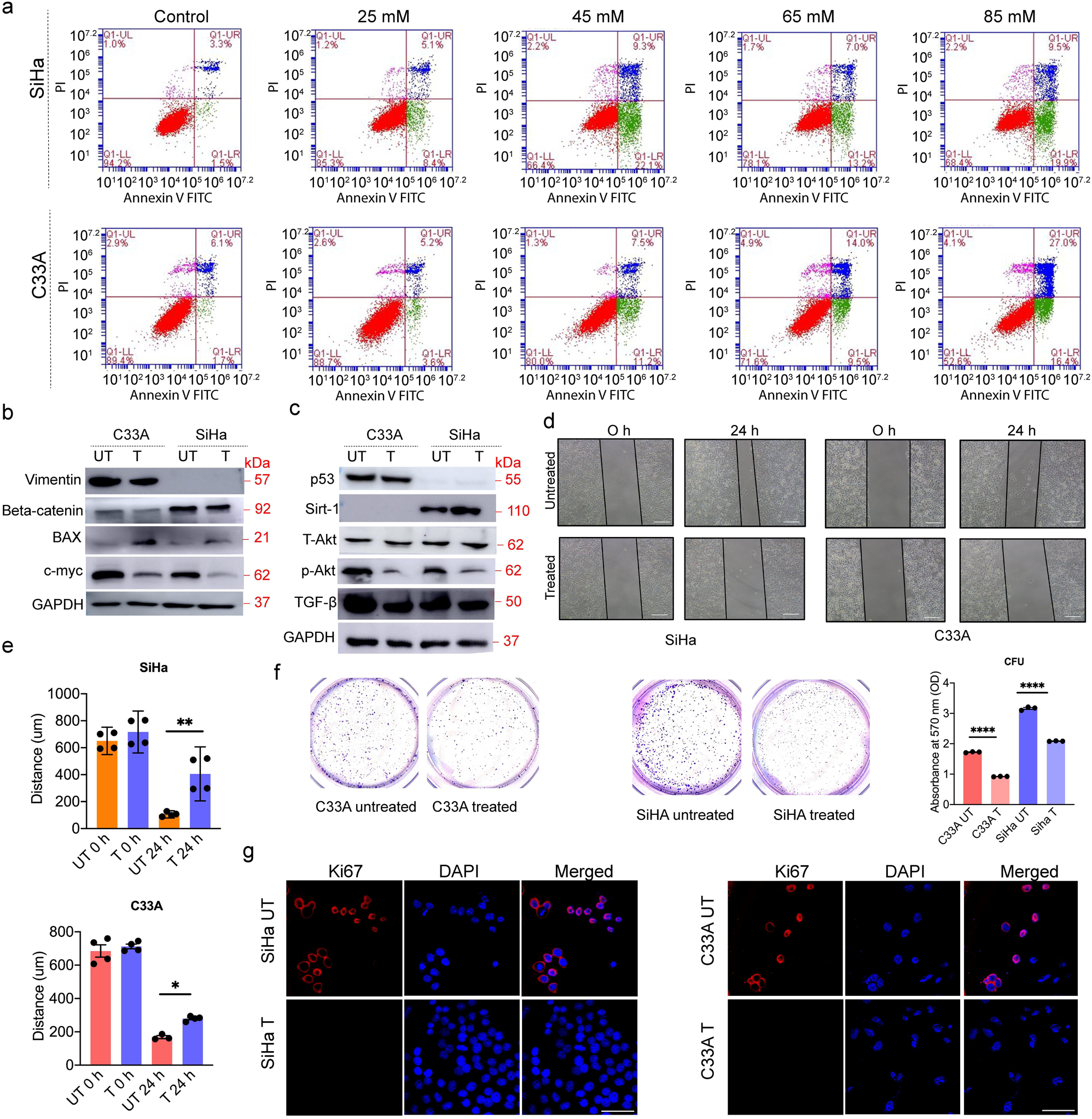
Therapeutic strategies towards cervical cancer pathogenesis. a. Each panel corresponds to a different treatment concentration (0-85 mM) with niacinamide for cervical cancer cells, SiHa (top row) and C33A (bottom row). The data points are color-coded to represent different cell populations or subpopulations. Red color represents potentially viable or actively proliferating cells, green represents late apoptotic, or necrotic state and blue color representing early apoptotic cells. With increase in the niacinamide concentration increase in the apoptotic cells were observed. The effect for HPV negative C33A cell line was greater than SiHa cell line. b. Western blot analysis of EMT (epithelial-mesenchymal transition) markers and apoptosis regulators. Expression levels of Vimentin, Beta-catenin, BAX (apoptosis regulator), and c-myc (oncogene) were measured in untreated (UT) and treated (T) C33A and SiHa cells. GAPDH serves as the loading control. c. Western blot analysis of key signaling proteins in C33A and SiHa cells. Levels of p53, SIRT1, total-Akt (T-Akt), phosphorylated Akt (p-Akt), and TGF-β were analyzed before and after treatment (UT vs. T). GAPDH serves as the loading control. d. Wound healing assay with cells in presence or absence of nicotinamide at different time points (0h and 24h) showing decrease in the proliferation of cells in presence of nicotinamide. d) The bar graph shows the relative distance of wound closure (μm) for SiHa and C33A cells under different treatment conditions (Untreated 0h, Untreated 24h, Treated 0h, and Treated 24h). e. Quantification of migration distances (in μm) over time (0 h, 12 h, 24 h) in SiHa and C33A cells under untreated and treated conditions. Significant reduction in migration is observed in treated cells. Data represent mean ± SEM. P-values: *P < 0.05; **P < 0.01. f. Image of colony-forming units (CFUs) determined by crystal violet staining. The number of viable colonies formed was more abundant in the untreated groups than in the treated groups for both the cell lines. Quantitative analysis of CFUs revealed significant reduction in colony formation nicotinamide treated cells (*p < 0.05). g. Immunofluorescence staining of proliferation marker Ki67 in untreated (UT) and treated (T) SiHa and C33A cells. DAPI (blue) is used as a nuclear counterstain. Reduced Ki67 staining (red) in treated cells indicates a decrease in cell proliferation. Scale bar is 50 μm.

Moreover, the changes in lipid metabolism observed in higher-grade tumors, including alterations in fatty acyls and phosphatidylcholines, suggest an increased reliance on fatty acid oxidation as an alternative energy source. This shift in metabolism, combined with changes in membrane lipid composition, supports the notion that metabolic reprogramming is a hallmark of cancer progression[31],[32]. The downregulation of enzymes such as PLPP2 and SMPD2 further emphasizes the role of lipid metabolism in regulating cellular signaling and membrane dynamics in cervical cancer.

The changes in niacinamide (a form of vitamin B3) and NAD^+^ levels suggest alterations in energy metabolism and redox homeostasis in cervical cancer cells. Niacinamide is a precursor for NAD^+^, which plays crucial roles in oxidative phosphorylation, glycolysis, and redox reactions[33]. NAM was significantly accumulated in Grade 2 cancer cells but not in Grade 3 cells (**Figure 2**). This difference might be due to the regulation of the salvage pathway that recycles NAM to produce NAD^+^, a crucial cofactor involved in energy metabolism, DNA repair, and cell signaling[34]. Cancer cells can meet their increased NAD^+^ demand through the de novo pathway (from tryptophan) or the more efficient salvage pathway[34]. In early tumor progression, cancer cells may rely more on the salvage pathway, leading to higher NAM levels due to upregulation of enzymes like NAMPT. However, as the tumor progresses to Grade 3, cancer cells may shift towards other pathways for NAD^+^ production, making the salvage pathway less important, resulting in lower NAM levels. Additionally, tumor microenvironment or epigenetic changes in Grade 3 cells may contribute to the differential regulation of the salvage pathway enzymes, affecting NAM levels. NAM has shown promising potential as a therapeutic agent for cervical cancer, particularly in targeting both HPV-positive and HPV-negative cell lines[35],[36]. Our findings also revealed that niacinamide (NAM) is involved in NAD^+^ metabolism, highlighting its potential as a therapeutic agent in cervical cancer. NAM treatment induced apoptosis in both HPV-positive (SiHa) and HPV-negative (C33A) cervical cancer cells. NAM is the amide form of niacin, an essential nutrient obtained from dietary sources and supplements. Numerous studies have demonstrated that NAM may be a potential candidate as an anticancer agent in many cancer cell models, such as pancreatic cancer[37], chronic lymphocytic leukemia[38], prostate carcinoma[39], and breast cancers[40],[41]. Therefore, in our present study we investigated the *in vitro* effect of NAM on cell proliferation and apoptosis in human cervical cancer cells to explore its potential mechanism. Interestingly, NAM’s effects on key EMT markers, such as vimentin and beta-catenin, were more pronounced in SiHa cells. The basal levels of vimentin differed between the two cell lines. Notably, NAM treatment resulted in downregulation of TGF-β in both cell lines. TGF-β is known to play a crucial role in promoting EMT and enhancing the metastatic potential of cancer cells. Therefore, the reduction of TGF-β could contribute to NAM’s ability to inhibit EMT and, consequently, metastasis. This underscores the significance of TGF-β in the context of NAM-mediated effects on cancer progression. This anti-metastatic effect was further supported by scratch assays showing impaired cell migration in NAM-treated cells.

The study also highlighted key differences in NAM’s mechanism of action between SiHa and C33A cells. qPCR analysis revealed a downregulation of CD38 in both cell lines following NAM treatment. Interestingly NAM treatment also upregulated SIRT1 (Sirtuin 1), an NAD^+^-dependent deacetylase in SiHa cells. It requires NAD^+^ as a cofactor for its enzymatic activity. NAM treated cells could increase NAD^+^ levels, potentially supporting SIRT1 activity leading to CD38 inhibition. qPCR analysis also revealed that upregulation of TFAM in SiHa cells on NAM treatment. This upregulation of Tfam may indicate enhanced mitochondrial biogenesis and oxidative metabolism, further supporting the notion that NAM could promote metabolic reprogramming in cancer cells. The increase in TFAM could be linked to the observed upregulation of SIRT1, which has been shown to play a role in mitochondrial function and biogenesis (**Figure 6f**). NAMPT was upregulated in SiHa but remained unchanged in C33A, indicating differential regulation of NAD^+^ metabolism. Additionally, PARP expression was downregulated in SiHa but upregulated in C33A, suggesting that NAM modulates NAD^+^ metabolism and DNA repair differently in these cell lines. The reduction of phosphorylated AKT (p-AKT) levels in both cell lines, while total AKT levels remained unchanged, and the downregulation of c-Myc further support NAM’s role in inhibiting pro-survival pathways and promoting apoptosis. Moreover, to evaluate the impact of NAM on cell proliferation, we examined Ki-67 expression, a well-established marker of cellular proliferation. NAM treatment led to a significant reduction in Ki-67 levels in both SiHa and C33A cells, indicating that NAM effectively inhibits cell proliferation in these cancer cell lines.

Thus, the study could provide valuable insights for developing targeted and personalized treatment strategies. Overall, our study provides a comprehensive view of the metabolic alterations in cervical cancer, offering potential therapeutic targets like NAM and pathways related to SIRT-1 and CD38 axis via the downregulation of EMT further supports the potential of NAM in inhibiting metastasis, making it a promising candidate for targeted therapy in cervical cancer.

## 4.0 Conclusion

Our study provides a detailed metabolic profile of cervical cancer, highlighting key alterations that accompany grade specific and HPV induced carcinogenesis. The identification of over 100 dysregulated metabolites in tumor tissues, including niacinamide, arginine, and various lipid metabolites, underscores the extensive metabolic reprogramming that supports cancer cell survival, rapid division, and adaptation to the tumor microenvironment. Specifically, changes in nucleic acid synthesis, energy metabolism, and redox balance were observed, along with alterations in lipid composition that reflect increased energy demands and structural needs in higher-grade tumors. The distinct metabolic shifts seen in our study not only offer potential biomarkers for early detection but also reveal promising therapeutic targets. Niacinamide (NAM) emerged as a significant regulator of NAD^+^ metabolism and oxidative stress management, with its role in reducing TGF-β and inhibiting EMT indicating its potential in hindering metastasis. The differential effects of NAM in HPV-positive and HPV-negative cell lines, such as the downregulation of CD38, upregulation of SIRT1, and modulation of EMT markers, further highlight its multifaceted role in cellular regulation. By elucidating the metabolic adaptations in cervical cancer, our findings support the potential of NAM and related pathways as candidates for targeted therapies. These insights into the SIRT1-CD38 axis, EMT inhibition, and differential NAD^+^ metabolism lay the groundwork for personalized therapeutic strategies, potentially enhancing the efficacy of treatments for cervical cancer. This study thus contributes to a growing understanding of how metabolic reprogramming shapes cancer progression and identifies metabolic interventions that could help curb cervical cancer proliferation and metastasis.

## 5.0 Material and Method

### 5.1. Chemicals and Reagents

All chemicals and reagents utilized in this study were of high purity and sourced from either Sigma-Aldrich (St. Louis, MO, USA) or Merck (Darmstadt, Germany). De-ionized and double-distilled water was procured from a Milli-Q (MQ) system (Millipore Corp., Bedford, MA, USA). The DNA extraction kit was acquired from Qiagen, USA. LCMS solvents and reagents were purchased from Honeywell (Charlotte, North Carolina, USA).

### 5.2. Human tumor biopsy sample procurement

A small cohort of freshly frozen human origin cervical cancer along with their corresponding non-cancer tissues of human origin were obtained from the National Tumor Tissue Repository, Tata Memorial Hospital, Mumbai, India. The study was approved by the Institutional ethics committee (IITB-IEC/2019/046), Indian Institute of Technology Bombay, Mumbai, India (IIT Bombay) and Institutional ethics committee (AUUP/IEC/APR/2022/6) of Amity University Uttar Pradesh, India. The National Tumor Tissue Repository situated at Tata Memorial Hospital, Mumbai India obtained informed consent from all tissue donors and all clinical investigation according to the principles expressed in the Declaration of Helsinki. The study involved 15 cervical cancer naïve tissues (11 tumor tissues and 4 adjacent non-cancerous tissues) between age 45-90 years. These cervical tissues were segregated grade wise as grade II and III as mentioned in Table S1. A minimum of four to five tissues per grade were selected for the study: however, in some grades. The sample size selection was based on several factors. In our cohort, we had only 4 number grade 2 samples and rest all were grade 3 samples. The table in the manuscript also shows a higher number of grade 3 cervical cancer samples compared to grade 2 samples. This reflects the tendency for cervical cancer to be detected more often at advanced stages, making earlier-stage samples rarer and more difficult to obtain in large numbers. Another key consideration in sample selection was the need to balance HPV positive and negative groups. The majority of the initial tissue samples were HPV positive, reflecting the high prevalence of HPV in cervical cancer cases. To create a more balanced comparison, we selected 11 cervical cancer samples that provided a relatively even distribution of HPV positive and negative cases. This careful selection allows for a more meaningful comparison between these two groups, despite the overall small sample size. All 11 cervical cancer samples used are from treatment-naive patients. These untreated samples are crucial for identifying potential early detection biomarkers, as they represent the natural molecular landscape of the disease before any therapeutic interventions. To address the limitations of the small sample size, we implemented rigorous statistical methods, extensive validation techniques, and also combine our data with publicly available dataset for meta-analysis.

### 5.3. H&E staining

The tissue sections underwent deparaffinization with a series of xylene washes, first with 100% xylene, followed by a mixture of xylene and ethanol in a 1:1 ratio. This was succeeded by rehydration with decreasing concentrations of ethanol from 100% to 50%, and a final wash with distilled water. Subsequently, the sections were then stained with 0.5% hematoxylin solution for 2 minutes, followed by a 1-minute staining with 0.8% eosin solution prepared in 95% ethanol. The slides were then incubated in xylene for 1 hour. Following this, the sections were mounted using dibutyl-phthalate polystyrene xylene (DPX) mounting medium and examined under a Leica DMi8 fluorescence microscope equipped with an Andor Zyla cCMOS camera (Oxford Instruments, UK).

### 5.4. DNA isolation and HPV status

Genomic DNA was extracted from cervical cancer tissue biopsies using QIA amp DNA Mini Kit (Qiagen) based on the manufacturer’s instructions. The genomic DNA quality and quantity were assessed using Nanodrop 2000 and Qubit (Thermo Fisher Scientific), respectively. The initial HPV detection was conducted as previously reported[42]. The genomic DNA from HPV positive cell lines such as SiHa was taken as control. PCR was carried out using MY09/MY11 primers (FP-5′ CGTCCMARRGGAWACTGATC-3′RP-5′ GCMCAGGGWCATAAYAATGC-3′) where M = A or C, W = A or T, Y = C or T and R = A or G. PCR was done using ready to use DreamTaq Green PCR Master Mix containing DNA Polymerase, buffer, MgCl_2_, and dNTPs (Thermo Scientific, USA). PCR condition is as follows-Initial denaturation at 94^ο^ C for 5 mins, followed by denaturation for 30 sec at 94^ο^ C, annealing at 55^ο^ C for 30 secs, extension at 72^ο^ C for 1 min for 35 cycles followed by final extension at 72^ο^ C for 10 min using Sure cycler 8800 PCR (Agilent Technologies USA). The PCR products were analyzed on 1.5% agarose gel electrophoresis to visualize the 450 bp amplicon.

### 5.4. Sample preparation for metabolomics data

#### 5.4.1 Tissue Metabolite Extraction

Approximately 30 mg of the tissue was weighed and cut into pieces using scalpel (surgical blade). Tissue pieces were added to bead beating tubes preloaded with 0.1 mm size glass beads (BioSpec Products, Cat. No. 11079101) of 250 µL in each tube to ensure equal volume in all tubes. Total volume of 450 µL methanol was added for the 30 mg of tissue (150 µL of solvent per 10 mg of tissue). Tissue lysis and metabolite extraction was then performed using bead beater (GENAXY Scientific Homogenizer, Model: GEN-Bioprep-6). The vibrating of the bead beater was 30 times/s for 40 s with 2 plus 2 cycles. The tubes were then centrifuged at 21,000g for 15 minutes at room temperature. The supernatant was collected, and the samples were dried in a vacuum concentrator and stored at −80 °C until the analysis. QC samples were prepared by pooling all the samples for normalization purposes.

#### 5.4.2. LC-MS/MS Method

Dried samples were reconstituted in 70 µL of 50% acetonitrile in water, vortexed, and filtered using a syringe (Phenex from Phenomenex) for analysis. Untargeted metabolomics was performed using Agilent 1290 Infinity II LC (Agilent Technologies Inc.) system that comprises a vacuum degasser, a temperature-controlled autosampler, a column compartment, and a quaternary pump. The HPLC system was coupled to the Agilent AdvanceBio 6545XT quadrupole-time-of-flight (Q-TOF) mass spectrometer along with a dual Agilent Jet Stream electrospray ionization source. Two different columns were used for the separation of metabolites namely Acquity BEH HILIC column (150[×[2.1[×[1.7[μm) (Waters Corporation) for polar metabolites and RPC18 (Reversed Phase) BEH C18 Acquity Premier column (150[×[2.1[×[1.7[μm) for separation of mid-polar and non-polar metabolites. Separate injections were done for positive and negative ionization modes and the data was acquired using MassHunter Software that was supplied with the instrument. LC gradient and MS method were taken from Dodia et al. 2024[43]. Data dependent acquisition (DDA) method, AutoMS/MS was then used for the data acquisition using the MassHunter acquisition software (Version 10.1.48, Agilent Technologies)

#### 5.4.3 Analysis of LC-MS/MS Data

The high-resolution LCMS data was analyzed using the MSOne Software Suite (available freely on https://msone.claritybiosystems.com), proprietary tool developed by Clarity Bio Systems India Pvt. Ltd. This software facilitated peak extraction with m/z and retention times (RT), and identification of metabolites. DDA mode of acquisition of samples facilitated the generation of fragment ion for several precursor ions. Metabolite annotation was then performed by matching fragment ions of precursor molecules against a reference library with MS2 data (Metabolomics Standards Initiative Level 2). The annotation names were standardized using RefMet or PubChem APIs. The data from the two columns and two ionization modes were separately filtered for blank subtraction, selecting the peaks only if the area of peak is atleast 5 times of the area in the blank (to exclude peaks present in blank), and imputed using LOD. To address instrument variability, QC samples were prepared by pooling individual samples, and the normalization technique Quality control–based robust LOESS signal correction (QC-RSLC) was applied[44]. A metabolite was considered significantly different between two cohorts if an unpaired t-test yielded a p-value<0.05 and the fold change > 1.5. FDR correction was not applied because of small sample size.

### 5.5. Analysis of transcriptomics dataset

To obtain an overview of the underlying genes affecting these metabolic pathways, we retrieved microarray data from NCBI (Gene Expression Omnibus: GSE63514) and performed correlational analysis with our metabolomics data. Genes that are commonly involved in metabolic reprograming of our respective enriched pathways were taken from GSEA. We observed that in accordance with our data, several genes affecting the major metabolic pathways were dysregulated when we overlapped the transcriptomics data with GSEA Human Gene Set data. Further we checked for the expression and survival analysis of those genes across the cervical cancer (CESC) patients via Box plot and Kaplan-Meier survival plot using GEPIA2.

### 5.6. Cell culture

C33A and SiHa were procured from the cell repository at the National Centre for Cell Science, Pune, India. C33A cells were also a kind gift from Dr. Amit Dutta and Dr. Alok C. Bharti. The cells were cultured in Minimum Essential Media Eagle (MEM) (Himedia, Mumbai, India) and Dulbecco’s modified Eagle’s medium (DMEM) (Himedia, Mumbai, India) respectively, supplemented with 10% FBS (Gibco Waltham, MA, USA) and 0.2% of Antibiotic-Antimycotic (Gibco Waltham, MA, USA). All cell cultures were kept at 37°C in a humidified chamber with 95% humidity and 5% CO_2_, using a CO_2_ incubator (Thermo Fisher Scientific).

### 5.7. Cell Viability assay

SiHa and C33A (10000 cells/well) were seeded in 96 well plate and were treated with different concentrations of nicotinamide for 24 hours ranging from 1 mM to 75 mM for both the cell lines. After 24 hours the media was discarded, and the cells were incubated with 100 μl crystal violet stain (0.5% crystal violet+20% methanol dissolved in distilled water) for 20 mins at 200 r.p.m in orbital shaker. The plate was then washed with tap water for destaining and left for drying overnight. 100 ul methanol was added in each well and incubated at orbital shaker for 20 mins at 200 r.p.m. Reading was taken at 570 nm using spectrophotometer and IC_50_ was calculated using Graphpad prism.

### 5.8. RNA isolation and Real time PCR

SiHa and C33A cells were cultured as mentioned above in presence and absence of NAM (40.1 mM NAM for C33A and 36.6 mM NAM for SiHa) for 24 hours. Untreated cells served as controls. After treatments, RNA was extracted using the TriZol method (Invitrogen) following the manufacturer’s instructions. RNA concentrations were determined with a NanoDrop spectrophotometer (Implen, West Village, CA, USA). RNA was reversed transcribed into cDNA (Complementary DNA) using iScript cDNA Synthesis kit (Bio-Rad) as per the recommended protocol. per the protocol. Quantitative real-time PCR (qRT-PCR) was then performed using the SYBR SYBR (PowerUP SYBR Green Master Mix; applied biosytems) on StepOne (applied biosystems) using Primers (**Supp Table 2**) on an Agilent MX3000P, with ROX as a passive dye, following the manufacturer’s guidelines.

### 5.9. Apoptosis Assay

Flow cytometry was employed to investigate the tumor-suppressive activities of niacinamide in SiHa and C33A cells. Both the cells were treated with different concentrations of niacinamide (NAM), 40.1 mM NAM for C33A and 36.6 mM NAM for SiHa respectively for 24[h to assess apoptosis. Cells, treated and untreated, were stained with Annexin V-AlexaFluor488 and PI using the Alexa Fluor 488 Annexin V/Dead Cell Apoptosis Kit (Invitrogen) as per the manufacturer’s protocol. Flow cytometry analysis was performed using an Accuri C6 flow cytometer (BD Biosciences), and the data were processed with BD Accuri C6 Software.

### 5.10. Western Blotting

Whole-cell lysates of SiHa and C33A cells in presence and absence of NAM (40.1 mM NAM for C33A and 36.6 mM NAM for SiHa) were obtained using RIPA buffer containing 150 mM NaCl, 0.5% sodium deoxycholate, 0.1% SDS, and 50 mM Tris-HCl at pH 8.0 (Sigma-Aldrich, #R0278). The total protein concentration in the lysates was determined using the Bradford assay. A protein amount of 50 µg from each sample was combined with a sample buffer containing 5% β-mercaptoethanol and heated at 100°C. The proteins were separated using 10% SDS-PAGE, followed by transfer onto a nitrocellulose membrane (Biorad) via the semi dry transfer method at for 25 minutes at a constant voltage of 16 V using a Bio-Rad transfer apparatus. After the transfer, the membrane was rinsed with TBST (TBS pH-7.6 with tween 20) and then blocked with 5% skimmed milk powder (dissolved in TBS) for 2 hours at room temperature on a rocker. Immunoblotting was performed with a primary mouse monoclonal antibody; anti-p53 (Santa Cruz Biotechnology DO-1; 1:500) or SIRT1 Polyclonal Antibody (BT-AP08324, Bioassay Technology Laboratory, 1:500, Japan), or Akt1 Polyclonal Antibody (BT-AP00347, Bioassay Technology Laboratory, 1:500, Japan), or Akt1(Phospho Ser129) Polyclonal Antibody (BT-AP06158, Bioassay Technology Laboratory, 1:500, Japan) or TGF beta 1 Rabbit pAb (A18692, Abclonal), Vimentin (D21H3) XP® Rabbit (5741, Cell Signaling Technology) or β-Catenin (D10A8) XP® Rabbit (8480, Cell Signaling Technology) or cMyc (D84C12, Cell Signaling Technology) overnight at 4°C. The blots were washed three times with TBST (0.1% Tween 20). Secondary immunoblotting was performed using secondary antibodies, mouse anti-GAPDH (1:2000; sc-365062, Santa Cruz Biotechnology, USA) or Anti-rabbit IgG, HRP-linked Antibody (7074, Cell Signaling Technology) (1:5000 dilution in 2% BSA in TBS) for 2 hours at room temperature with continuous rocking. Following this, the blots were washed three with TBST (0.1% Tween 20) and treated with Clarity Western ECL Substrate (Bio-Rad, USA, #1705060). Visualization was achieved using the ImageQuant LAS 500 chemiluminescence ChemiDoc system coupled with a CCD camera (Cytiva, USA). Three independent experiments were conducted.

### 5.11. Wound healing assay

SiHa and C33A cells were grown in 10% DMEM and MEM media respectively as mentioned above in presence and absence of NAM (40.1 mM NAM for C33A and 36.6 mM NAM for SiHa) following the method by Rodriguez et al. (2005)[45] . The cells were plated in a 6-well plate, while untreated cells served as controls. After rinsing with PBS (pH 7.4), a scratch was made with a sterile plastic tip. Detached cells were removed by washing with PBS, and the cells were then exposed to medium containing mitomycin C (0.5 μg/ml, Sigma-Aldrich, St. Louis, MO, USA) to inhibit cell proliferation, ensuring accurate measurement of migration. Cells were incubated for an additional 24 hours to allow for wound closure. Images were captured at the initial and final time points using a Leica DMi1 bright-field microscope at 10x magnification, and wound closure was analyzed using ImageJ software. The cell migration rate (μm/h) was calculated using the formula: cell migration rate = [(initial wound width - final wound width) / time taken (hours)], based on three independent experiments.

### 5.12. Colony forming assay

SiHa and C33A cells (1000 cells per well) were seeded in 60 mm dishes and cultured in presence and absence of NAM (40.1 mM NAM for C33A and 36.6 mM NAM for SiHa). The single colonies were then allowed to grow for 14 days, fresh media was added every 3^rd^ day after giving PBS wash. Afterwards the cells were stained with crystal violet stain then washed with tap water for destaining and left for drying overnight. 1ml of methanol was added and incubated at orbital shaker for 20 mins at 200 r.p.m[46]. Reading was taken at 570 nm using spectrophotometer.

### 5.13. Immunofluorescence

SiHa and C33A cells were cultured as mentioned above in presence and absence of NAM (40.1 mM NAM for C33A and 36.6 mM NAM for SiHa). Cells were fixed onto coverslips with 4% paraformaldehyde (Himedia, Mumbai, India) for 20 minutes at room temperature (25°C). Fixed cells were rinsed with PBS (pH 7.4) and permeabilized with 0.2% Triton X-100 (Sigma-Aldrich, St. Louis, MO, USA) in PBS for 10 minutes. After another PBS wash, the cells were blocked with 5% bovine serum albumin (BSA, Himedia, Mumbai, India) in PBS for 1 hour to prevent non-specific binding. The cells were then incubated at 4°C overnight with a 1:200 dilution of mouse monoclonal IgG ki67 primary antibody (sc-23900, Santa Cruz Biotechnology, Dallas, TX, USA). After incubation, cells were rinsed three times with PBST (PBS containing 0.1% Tween 20, pH 7.4). Coverslips were then treated with a 1:500 dilution of Alexa Fluor 555-conjugated goat anti-mouse IgG secondary antibody (A-21428, Invitrogen, USA) for 2 hours at 25°C. Following three washes with PBST, the cells were further incubated with DAPI(0.1ug/ml) for 15 mins. The coverslips were mounted on slides using a medium containing 1% 1,4-diazabicyclo[2.2.2]octane (DABCO, Sigma-Aldrich, St. Louis, MO, USA) in a 90% glycerol/10% PBS solution. The coverslips were observed using a Leica TCS SP8 confocal microscope and images were analyzed using ImageJ software.

### 5.14. Statistical analysis

The statistical significance was calculated in this study and assessed through one-way ANOVA followed by Tukey’s multiple comparison test. Significance levels were denoted as *P<0.05, **P<0.01, and ***P<0.001, with non-significant results marked as NS (P≥0.05). GraphPad Prism was employed for statistical analysis.

### 5.15. Data availability

Metabolomics data is deposited to the EMBL-EBI MetaboLights database (DOI: 10.1093/nar/gkad1045, PMID:37971328) with the identifier MTBLS10405. Once the status of your submission is updated as ‘Public’, the complete dataset can be accessed here https://www.ebi.ac.uk/metabolights/MTBLS10405

## Acknowledgment and Funding

The authors would like to acknowledge the National Tumor Tissue Repository (NTTR), Indian Council for Medical Research (ICMR) at the Tata Memorial Hospital (TMH), Mumbai, India, for providing human cervical cancer and normal tissue. The authors thank Central Instrumentation Facility of University of Delhi South Campus for confocal microscope facility. The Flow cytometry related data acquisition was performed with help of Manoj Gupta, Department of Science and Technology (DST) and FIST sponsored (SR/FST/LS-II/2017/115) flow cytometry facility at AIMMSCR, Amity University, Noida. The authors would like to acknowledge Dr. Amit Dutt and Dr. Alok C. Bharti for their kind gift of C33A cells. The author would also like to thank Prof. Bhudev C. Das for his generous gift of SiHa cells. The authors would like to acknowledge the DBT/Wellcome Trust India Alliance Fellowship [IA/E/17/1/503663] awarded to Shinjinee Sengupta and DST SERB (CRG/2023/001776) for financial support.

