## Supplementary File for "Metabolomic profiling reveals grade-specific niacinamide accumulation and its therapeutic Potential via SIRT1-CD38-EMT axis modulation in cervical cancer progression"

The supplementary File contains

4 Figures
1 Table

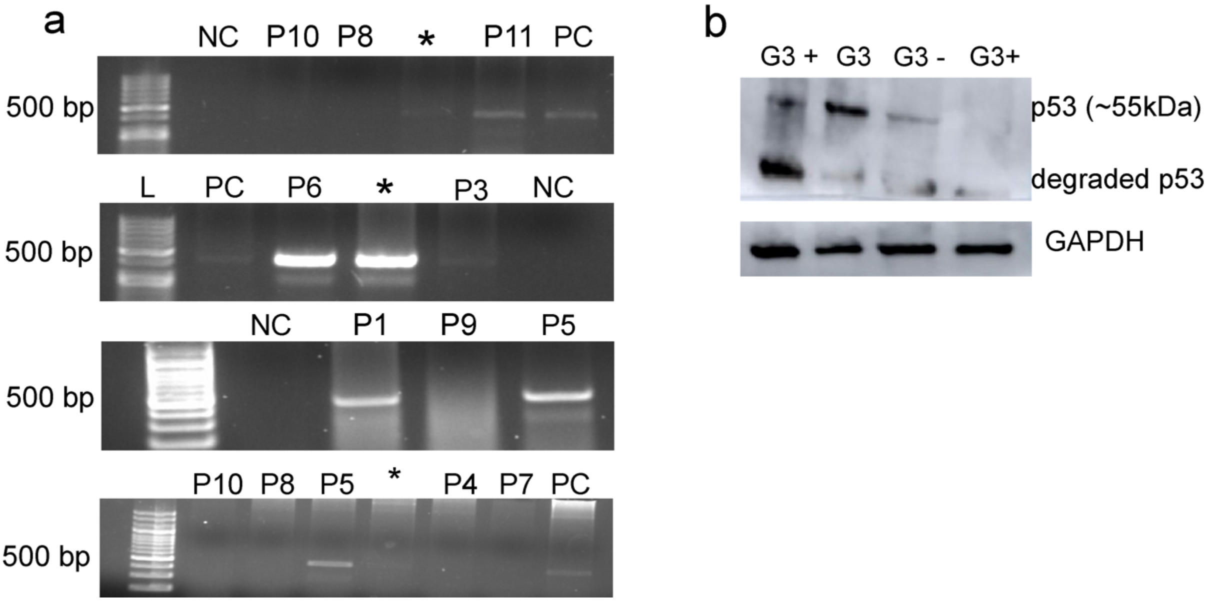


**Supplementary Figure 1. Characterization of the cervical tumor biopsy cohort.** a. Agarose gel images showing HIV bands when PCR was performed with L1 structural primers to detect the presence of HIV. The lanes marked as * refer to the samples that were not studied using metabolomics and were excluded. NC refers to negative control while PC refers to positive control. b. Western blot with anti-p53 antibody to study the expression level of p53 in the tissue cohort. G3 refers to Grade 3 cervical cancer tissues. "+" and "-" refers to the presence or absence of HPV infection respectively.

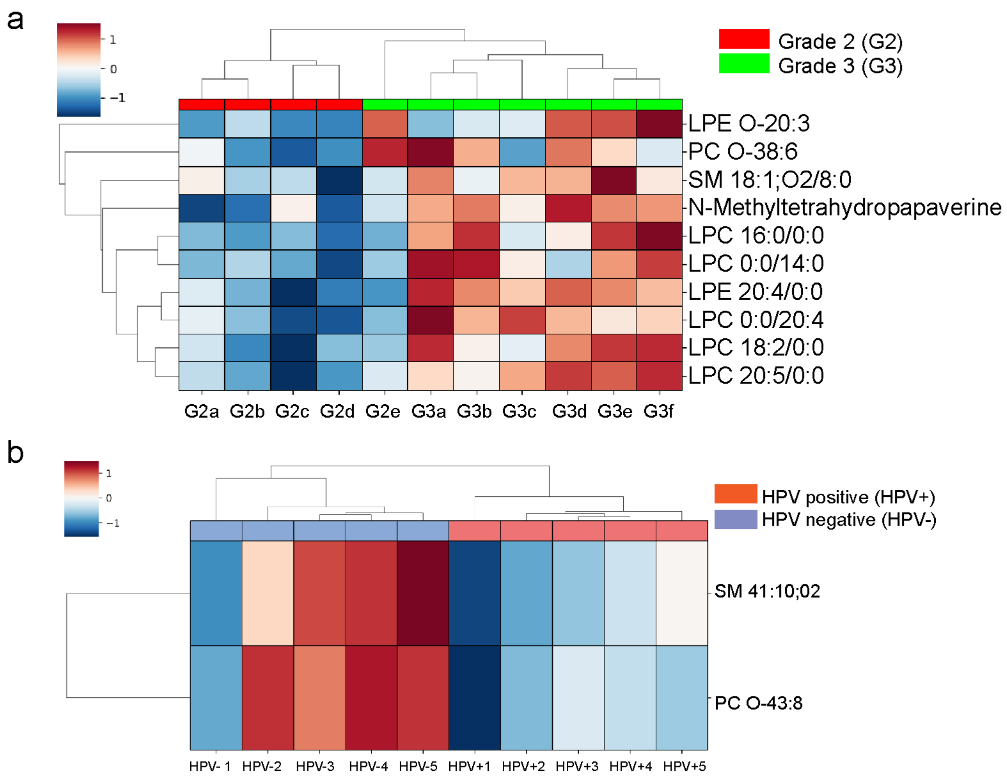


**Supplementary Figure 2. Metabolic profiling of Grade 2 vs. Grade 3 tumors and HPV-positive vs. HPV-negative samples.** (a) Heatmap showing hierarchical clustering of lipid species identified in Grade 2 (G2, red) and Grade 3 (G3, green) tumor samples. Blue represents lower levels of the lipid, while red represents higher levels, with a color scale from -1 to 1 indicating the intensity. The lipids included are lysophosphatidylethanolamines (LPE), lysophosphatidylcholines (LPC), phosphatidylcholines (PC), and sphingomyelins (SM). Lipid classes such as LPE O-20:3, LPC 16:0/0:0, and PC O-38:6 demonstrate differential expression between tumor grades. (b) Heatmap showing hierarchical clustering of lipid species in HPV-positive (orange) and HPV-negative (blue) samples. Samples are grouped based on HPV status, and the heatmap represents differences in lipid profiles, with red indicating higher lipid expression and blue indicating lower expression. Notable lipid species include SM 41:10/02 and PC O-43:8, which exhibit differential expression in relation to HPV status.

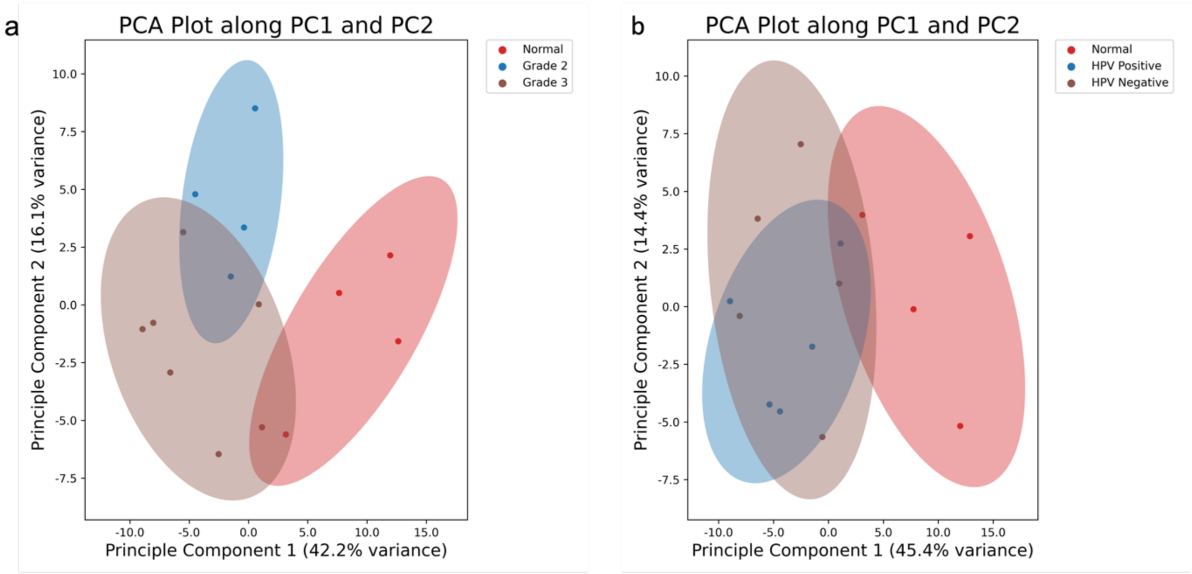


**Supplementary Figure 3. Principal Component Analysis (PCA) of Samples Based on Histological Grading and HPV Status.** a. PCA plot showing the separation of samples based on histological grading. Samples are classified as Normal (red), Grade 2 (blue), and Grade 3 (brown). PC1 and PC2 explain 42.2% and 16.1% of the variance, respectively. Clear segregation is observed among the different histological grades. b. PCA plot depicting the separation of samples based on HPV status. Samples are grouped as Normal (red), HPV Positive (blue), and HPV Negative (brown). PC1 and PC2 account for 45.4% and 14.4% of the variance, respectively. The plot highlights the distribution of HPV status across the samples, with partial overlap between the HPV positive and negative groups.

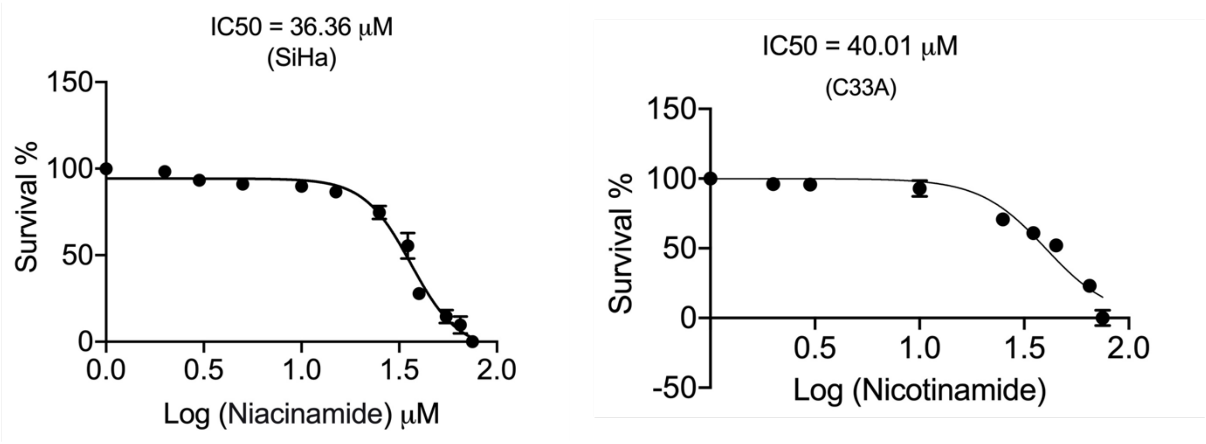


**Supplementary Figure 4. Calculation of $IC_{50}$ of Cervical cell lines.** a. Sigmoidal curve from crystal violet assay indicating the $IC_{50}$ value on HPV positive Siha (left) and HPV negative C33A (Right) cells. Each data point is an average derived from three independent experiments.

**Supplementary Table 1. List of cervical cancer tumor biopsies**

| S. No. | Patient Number | Cervical tissue Type | HPV status | Age | Stage | Treatment | Diagnosis |
|---|---|---|---|---|---|---|---|
| P1 | 6103 | Grade 2 | Positive | 56 | T1N0M0 | Naive | Moderately differentiated squamous carcinoma |
| P2 | 8664 | Grade 2 | NA* | 68 | T1BN0M0 | Naive | Moderately differentiated squamous carcinoma |
| P3 | 6470 | Grade 2 | Positive | 52 | T1B2N0M0 | Naive | Moderately differentiated non keratinising squamous carcinoma |
| P4 | 3317 | Grade 2 | Negative | 49 | T1N0M0 | Naive | Invasive moderately differentiated squamous carcinoma |
| P5 | 1248 | Grade 3 | Positive | 58 | T1N0M0 | Naive | Poorly differentiated squamous carcinoma |
| P6 | 2284 | Grade 3 | Positive | 53 | T1N0M0 | Naive | Poorly differentiated squamous carcinoma |
| P7 | 6657 | Grade 3 | Negative | 63 | T2N0M0 | Naive | Poorly differentiated squamous carcinoma |
| P8 | 1016 | Grade 3 | Negative | 85 | T2N1M0 | Naive | Poorly differentiated squamous carcinoma |

| | | | | | | | |
|---|---|---|---|---|---|---|---|
| P9 | 1721 | Grade 3 | Negative | 64 | T1N1M0 | Naive | Poorly differentiated squamous carcinoma |
| P10 | 5748 | Grade 3 | Negative | 61 | T1N0Mx | Naive | Poorly differentiated squamous carcinoma |
| P11 | 6933 | Grade 3 | Positive | 64 | T2N1M0 | Naive | Poorly differentiated squamous carcinoma |
| P12 | 6933 | Normal | NA | | | | |
| P13 | 72N | Normal | NA | | | | |
| P14 | 1721 | Normal | NA | | | | |
| P15 | 0311N | Normal | NA | | | | |

Patient details along with grade and HPV status

*NA- HPV status is unavailable due to sample limitation

**Supplementary Table 2. List of primers used in the study.**

| S. No | Name | Sequence |
|---|---|---|
| 1 | SIRT-1 | Forward-TAGACACGCTGGAACAGGTTGC |
| | | Reverse-CTCCTCGTACAGCTTCACAGTC |
| 2 | Tfam | Forward- GTGGTTTTCATCTGTCTTGGCAAG |
| | | Reverse- TTCCCTCCAACGCTGGGCAATT |
| 3 | CD38 | Forward-TCTTGCCCAGACTGGAGAAAGG |
| | | Reverse- TGGACCACATCACAGGCAGCTT |
| 4 | NAMPT | Forward- AGGGTTACAAGTTGCTGCCACC |
| | | Reverse- CTCCACCAGAACCGAAGGCAAT |
| 5 | PARP | Forward- CCAAGCCAGTTCAGGACCTCAT |
| | | Reverse- GGATCTGCCTTTTGCTCAGCTTC |
